# Serine peptidases and increased amounts of soluble proteins contribute to heat priming of the plant pathogenic fungus *Botrytis cinerea*

**DOI:** 10.1101/2023.03.23.534054

**Authors:** Mingzhe Zhang, Naomi Kagan Trushina, Tabea Lang, Matthias Hahn, Metsada Pasmanik Chor, Amir Sharon

## Abstract

*Botrytis cinerea* causes grey mold disease in leading crop plants. The disease develops only at cool temperatures, but the fungus remains viable in warm climates and can survive periods of extreme heat. We discovered a strong heat priming effect in which the exposure of *B. cinerea* to moderately high temperatures greatly improves its ability to cope with subsequent, potentially lethal temperature conditions. We showed that priming promotes protein solubility during heat stress and discovered a group of priming-induced serine-type peptidases. Several lines of evidence, including transcriptomics, proteomics, pharmacology, and mutagenesis data, link these peptidases to the *B. cinerea* priming response, highlighting their important roles in regulating priming-mediated heat adaptation. By imposing a series of sub-lethal temperature pulses that subverted the priming effect, we managed to eliminate the fungus and prevent disease development, demonstrating the potential for developing temperature-based plant protection methods by targeting the fungal heat priming response.

**Importance:** Priming is a general and important stress adaptation mechanism. Our work highlights the importance of priming in fungal heat adaptation, reveals novel regulators and aspects of heat adaptation mechanisms, and demonstrates the potential of affecting microorganisms, including pathogens through manipulations of the heat adaptation response.

## Introduction

*Botrytis cinerea* is a notorious plant pathogen that causes grey mold disease, leading to massive crop losses worldwide. Its optimum growth temperature is 18–22°C, and grey mold disease is widespread in relatively cool environments. At temperatures just a few degrees higher than the optimum, the disease completely disappears; however, the fungus remains viable in warm climate and resumes growth and infection when temperatures drop to the optimal range. How *B. cinerea* copes with high temperatures and remains viable in warm environments, including occasional extreme heat waves, is unclear.

Temperature shifts alter the expression of thousands of genes. In budding yeast, the most significantly enriched gene ontology (GO) categories that are upregulated under heat stress include heat shock proteins (HSPs), oligosaccharide metabolism, protein folding, and protein catabolic and recycling processes (1, 2). Proteins in these categories, as well as the disaccharide trehalose, help maintain protein homeostasis (proteostasis) (3, 4). Downregulated genes are typically enriched in translation and ribosome biogenesis, which correlates with growth arrest at high temperatures (1, 5). Apart from chaperones, the roles of most heat-induced genes and their protein products remain enigmatic. Most of these genes might help replenish functional proteins that are lost due to increased heat-induced protein aggregation and degradation (2). Additional studies have highlighted the central roles of proteostasis and protein aggregation in heat adaptation (6–11).

Excess heat causes protein misfolding and aggregation, rapidly endangering cell viability. To cope with proteotoxic damage, organisms have evolved a quality control (QC) system that removes misfolded proteins by promoting their refolding using chaperones or by proteolytic degradation through the ubiquitin–proteasome and autophagy systems (6, 12). When the stress level exceeds the capacity of these systems, proteins form aggregates, new protein translation slows and growth is arrested (2, 10, 13–15). The sequestration of misfolded proteins into insoluble aggregates is reversible and likely cytoprotective, as it removes defective and misfolded toxic proteins from the soluble phase (8, 9). When heat stress is relieved, protein aggregates are either disaggregated with the aid of the Hsp70, Hsp40, and Hsp110 chaperone network (4, 10, 11, 13, 16) or eliminated by the proteasome and selective autophagy (6, 17).

Although the protein QC system is evolutionarily conserved, fungi exhibit fundamental differences in their responses to moderate and severe heat stress. The upregulation of molecular chaperones and certain metabolic processes represents the predominant response of yeast to moderately high temperatures (MHTs; 37°C), whereas under severely high temperatures (SHTs; 42-46°C), protein aggregates form and translation shuts down (2, 13). Similarly, Gibney et al. (2013) (18) showed that the specific genes that mediate the responses to severe and moderate heat stress differ considerably, and suggested that the survival of yeast cells after heat shock might depend on relatively few genes. Importantly, pre-exposure to mild heat stress can mitigate heat damage while preparing the organism for more severe, potentially lethal stress, a phenomenon known as priming or acquired stress resistance (19–22). This general mechanism, which is well known in plants, is also widespread in microorganisms, helping them cope with changing conditions (21, 23–27). Studies in budding yeast showed that priming is dependent on nascent protein synthesis during moderate but not severe heat stress (18, 20). In filamentous fungi, priming can boost stress tolerance, however not all fungi respond in the same manner, and the specific mechanisms may differ depending on the fungal lifestyle (23, 27, 28).

Here we show that MHT had a strong priming effect on *B. cinerea*, namely it greatly improved the ability of the fungus to cope with a subsequent potentially lethal heat stress. Using transcription profiling and proteomics analysis of soluble and aggregated proteins, we were able to differentiate the priming from a general heat stress effect and showed that priming improves protein solubility during severe heat stress. Among priming-associated proteins, we identified one serine-type peptidase that to be essential for the priming response.

## Results

### *B. cinerea* growth varies substantially with temperature

We characterized spore germination and the growth of germ tubes (GTs) and mycelia at selected temperatures between 22°C and 37°C. Both spore germination and mycelial growth decreased with increasing temperatures above 22°C (Fig. S1a and S1b). Spores were less sensitive to supra-optimal temperatures than mycelia, as spore germination was completely blocked at 37°C, whereas mycelial growth was completely blocked at 32°C. By contrast, GT elongation increased at 22–29°C and then declined sharply (Fig. S1c). Spore germination was delayed at 29°C compared to 22°C, whereas the GT growth rate was much faster at this temperature, resulting in ∼50% longer GTs after 12 h of incubation at 29°C vs. 22°C (Fig. S1d). Exposure of GTs to oxidative (H_2_O_2_) and cell wall (SDS) stress generated similar response curves (Fig. S1e and S1f), suggesting that the accelerated GT growth at MHTs represents a general response to moderate stress. Based on these results, we defined 29°C as MHT and 37°C as SHT, as incubation at 29°C induced stress-adaptive developmental changes, whereas germination and growth were completely blocked at 37°C. We also determined that the early GT (8–12 h) is a suitable material in which to study the effects of MHTs and SHTs.

### MHTs prime cells to better cope with SHTs

We produced GTs at 22°C or 29°C, transferred them to 37°C and monitored cell death by propidium iodide (PI) staining. Transfer from 22°C to 37°C resulted in massive cell death, with ∼80% of GTs showing positive PI staining after 6 h and 100% after 12 h at 37°C (Fig. 1a and S2a). When GTs were produced at 29°C, the average rate of cell death following transfer to 37°C was drastically lower, with <5% PI-positive cells after 6 h and <40% after 12 h at 37°C. Similarly, membrane potential was severely compromised in GTs that were transferred from 22°C to 37°C, but not in those transferred from 29°C to 37°C (Fig. 1b and S2b). Next, we tested the effect of incubation at 29°C on recovery of the fungus after exposure to 37°C. Colonies originating from GTs produced at 22°C and transferred to 37°C had smaller diameters and less dense mycelia than control colonies kept at 22°C for 3 d, whereas GTs produced at 29°C developed colonies similar in diameter to control colonies (Fig. 1c and S2c). Therefore, preincubation at 29°C (MHT) has a strong priming effect that enabled cells to better cope with successive exposure to the potentially lethal SHT of 37°C.

**Fig 1.**
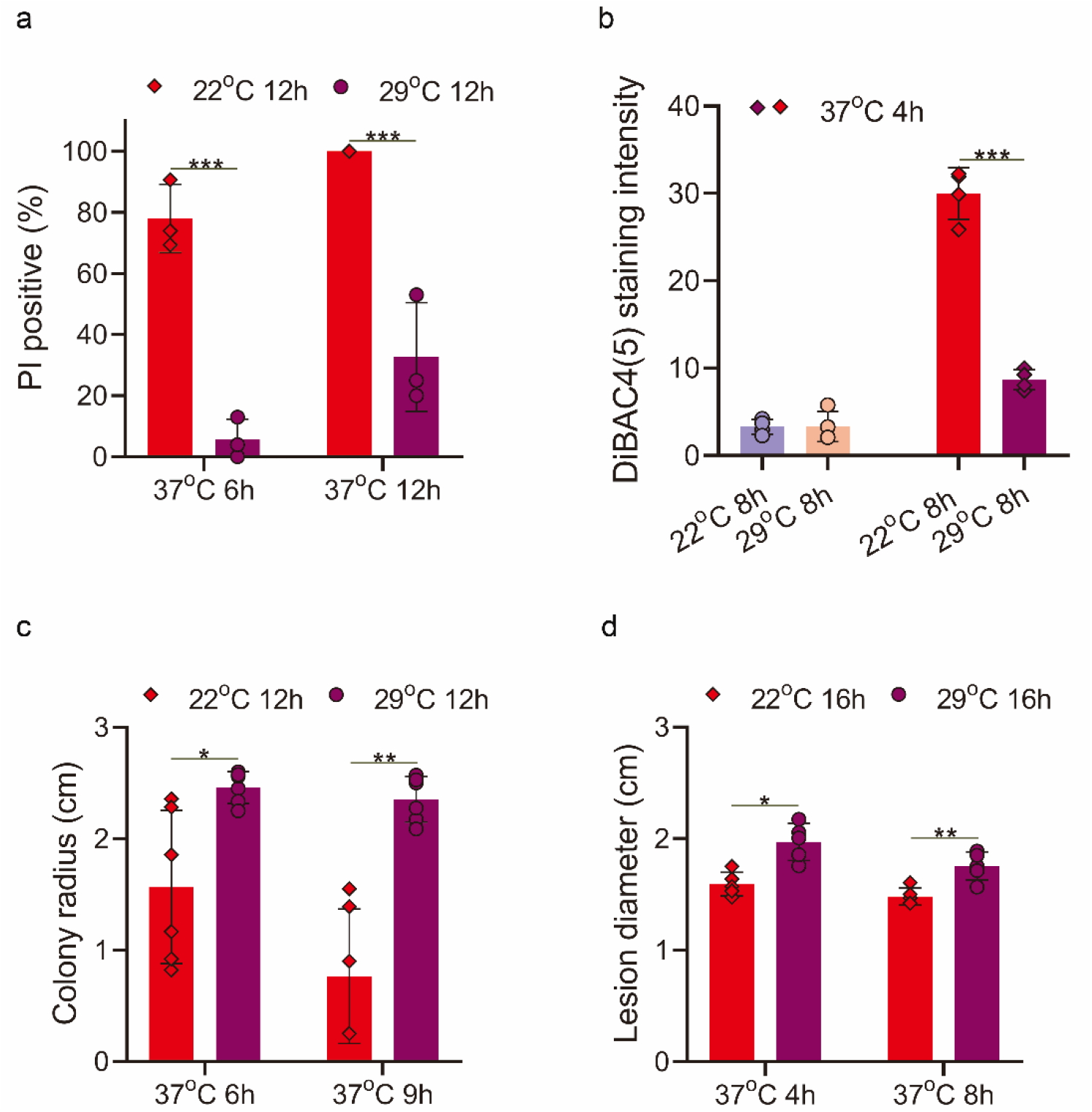
Pre-exposure to moderately high temperatures improves heat tolerance in *B. cinerea*. **a**. Measurement of cell death by PI staining. GTs were produced at 22°C or 29°C, transferred to 37°C for 6 or 12 h and stained with PI, and the percentage of PI-positive (dead) GTs determined. **b**. Measurement of cell membrane potential. GTs were produced at 22°C or 29°C, transferred to 37°C for 4 h and stained with DiBAC4(5). Images were captured under a fluorescence microscope, and the cell membrane potential of each GT was quantified by measuring fluorescence intensity using ImageJ. Higher fluorescence intensity represents greater disruption of the membrane potential. **c**. Recovery of colony growth after exposure to severe high temperatures with or without priming. GTs were produced at 22°C (no priming) or 29°C (priming), transferred to 37°C for 6 or 9 h and moved back to 22°C. Colony diameter was recorded after 3 d at 22°C. **d**. Effect of priming on pathogenicity. Inoculated plants were incubated for 16 h at 22°C or 29°C, transferred to 3f7°C for 4 or 8 h and incubated at 22°C for 3 d, and the lesion diameter was recorded. Graphs represent three (**a**), four (**b**), five (**d**) and six (**c**) biological replications with overlaid individual data points. Values are presented as the mean of replicates ± s.d. Statistical differences were determined according to unpaired two-tailed Student’ t-test (\**P* < 0.05; \*\**P* < 0.01; \*\*\**P* < 0.001).

### Temperature influences fungal viability and pathogenicity

We inoculated the leaves of French bean (*Phaseolus vulgaris*) plants with spore suspension, incubated them at 22°C or 29°C for 16 h, transferred them to 37°C for 4 h and then incubated them at 22°C for 3 d. When inoculated plants were incubated at 29°C followed by 37°C, they developed less severe symptoms than plants incubated at 22°C throughout the experiment (control) but more severe symptoms than those incubated at 22°C + 37°C, with average lesion diameters of 2.0 cm, 2.75 cm and 1.6 cm, respectively (Fig. 1d and S2d). Extended incubation for 8 h at 37°C had only mild effects on disease symptoms, with slightly reduced lesion size under both treatments.

Exposure of the fungus to 37°C for 6 h without acclimation at 29°C led to cell death, but it did not completely kill the fungus and did not prevent disease development. Spores were much less sensitive to heat than GTs: only a small fraction were killed after a lengthy incubation at 37°C (Fig. 2a), and even at 42°C, it took >10 h to kill ∼100% of spores (Fig. 2b). We hypothesized that a series of short pulses of SHT separated by periods of optimal temperature (OT) might bypass the priming effect and help build up lethal levels of cellular damage(1), with little or no negative effect on the plant. To test this hypothesis, we exposed GTs to 42°C for 2 h, followed by 22 h at 22°C. Less than 10% of the cells died after one cycle. However, two cycles of {42°C (2 h) / 22°C (22 h)} resulted in ∼85% cell death, and 100% of the GTs were killed by three cycles of treatment (Fig. 2c and 2d). To evaluate the effects of heat treatments on pathogenicity, we inoculated plants and exposed them to 42°C for 2 h, followed by 22°C. Disease was completely prevented by three cycles of {42°C (2 h) / 22°C (22 h)} treatment, and the plants showed no visible stress symptoms (Fig. 2e).

**Fig 2.**
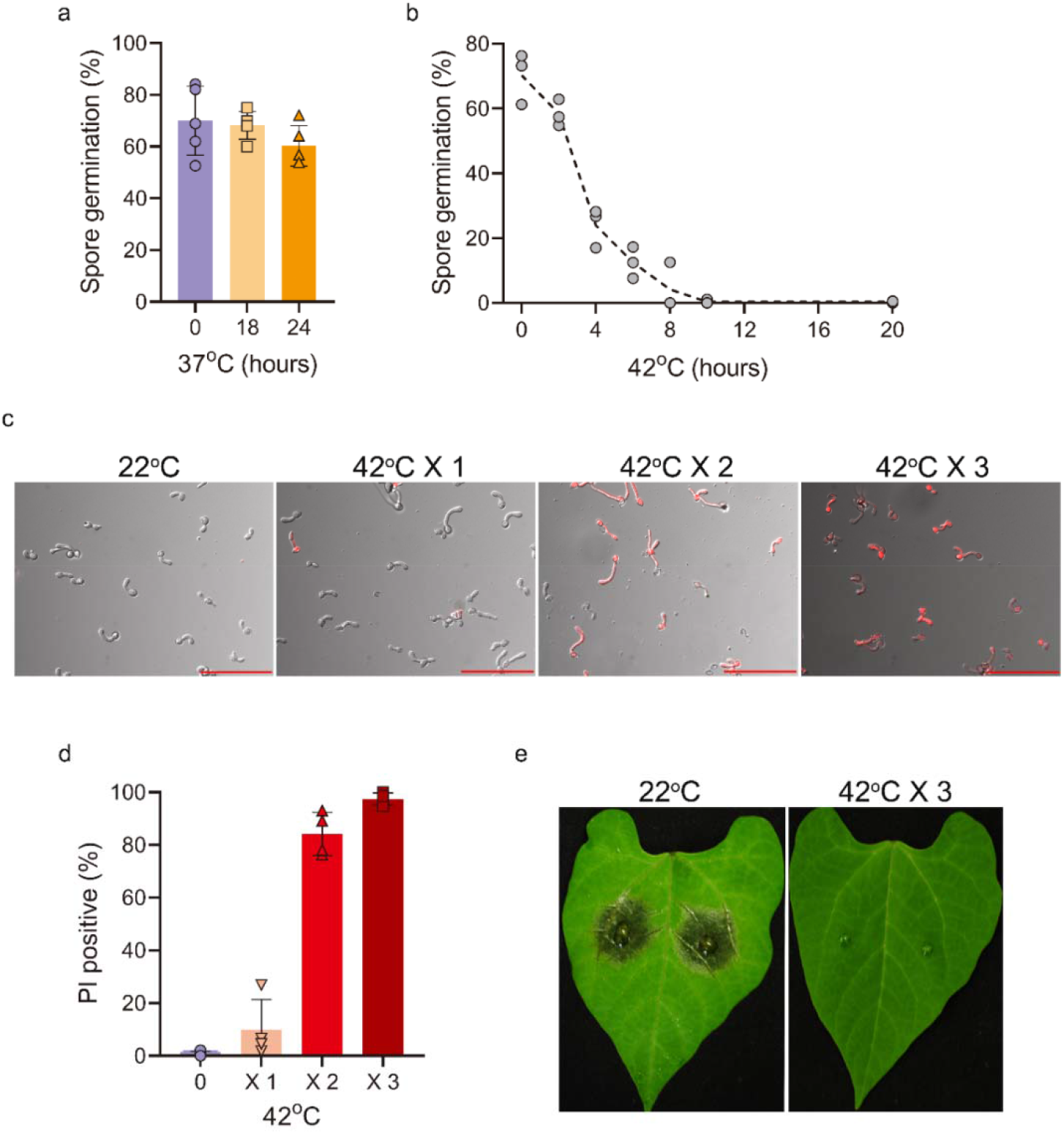
Effects of different heat regimes on fungal viability and pathogenicity. **a**. Spore germination after exposure to 37°C. Cultures were produced on PDA, seven-day-old cultures were exposed to 37°C for 18 or 24 h, spores were collected and incubated at 22°C for 3 d, and germination rates were determined. **b**. Cultures were treated at 42°C, and spores were collected at different time intervals and incubated at 22°C for 3 d and germination rates determined. **c**. Combined DIC and fluorescence (rhodamine filter) microscopic images of GTs following exposure to 42°C and staining with PI. Scale bars, 100 μm. **d**. Increased cell death following repetitive cycles of exposure to 42°C and recovery at 22°C. Spores were germinated at 22°C for 6 h, and the GTs were transferred to 42°C for 2 h and then 22°C for 22 h. The cycle was repeated three times. GTs were stained with PI after each cycle, and the levels of cell death were determined. **e**. Effects of {42°C (2 h)/22°C (22 h)} cycles on pathogenicity. Inoculated *P. vulgaris* plants were incubated for 6 h at 22°C, transferred to 42°C for 2 h and incubated at 22°C for 22 h, and the cycle was repeated three times. Photographs were taken after the third cycle. Control plants were maintained at 22°C for the entire duration of the experiment. Graphs represent three (**b**), four (**c**) and five (**a**) biological replications with overlaid individual data points. For **a** and **d**, values are presented as the mean of replicates ± s.d.

### Search for regulators of the heat priming response

To unravel the molecular mechanisms behind the priming response, we aimed to identify genes and proteins associated with priming induction and execution.

#### RNA-seq analysis

We grew fungi as outlined in Table S1 with four replications per treatment and extracted and sequenced RNA from the samples. We identified 7,993 differentially expressed genes (DEGs), including 4,814 with a positive false discovery rate (pFDR) < 0.05 and fold change (lFCl) ≥ 2. Of these, 757 DEGs were detected under MHT, 3,811 under SHT-P (priming conditions) and 3,409 under SHT (Fig. S3d and Table S3). GO enrichment analysis indicated that the main functional GO categories (Table S4) of the genes downregulated under SHT-P were ribosome function (GO:0005840, GO:0003735, GO:0015935) and translation (GO:0006412, GO:0005852, GO:0003743), which is in line with the finding that the translation machinery shuts down under severe heat stress (2). Two GO categories were specifically downregulated under SHT: zinc ion binding (GO:0008270) and telomere maintenance (GO:0000723). Genes in these categories trigger cell cycle arrest and cell death in response to cellular stress (29–31). Among the upregulated genes, only two functional categories were shared between MHT and SHT-P: proteolysis (GO:0006508) and serine-type peptidase (GO:0008236).

To identify the most significant DEGs, stricter selection (pFDR < 0.05, lFCl ≥5) were applied, and 1,329 DGEs were determined, 144 under MHT (29°C), 878 under SHT (22°C + 37°C) and 965 under SHT-P (Fig. S3e). Heatmap analysis separated the data into two main clusters: OT & MHT and SHT & SHT-P (Fig. 3a). Deeper inspection revealed two small gene clusters that we interpreted as priming-related genes: cluster A, comprising 39 genes that were upregulated at MHT but not OT or SHT; and cluster B, including 16 of the 39 genes (cluster A) that were exclusively upregulated at both MHT and SHT-P (Table S5). Four serine peptidase-encoding genes were shared between the two clusters, and two more were specific to cluster A.

**Fig 3.**
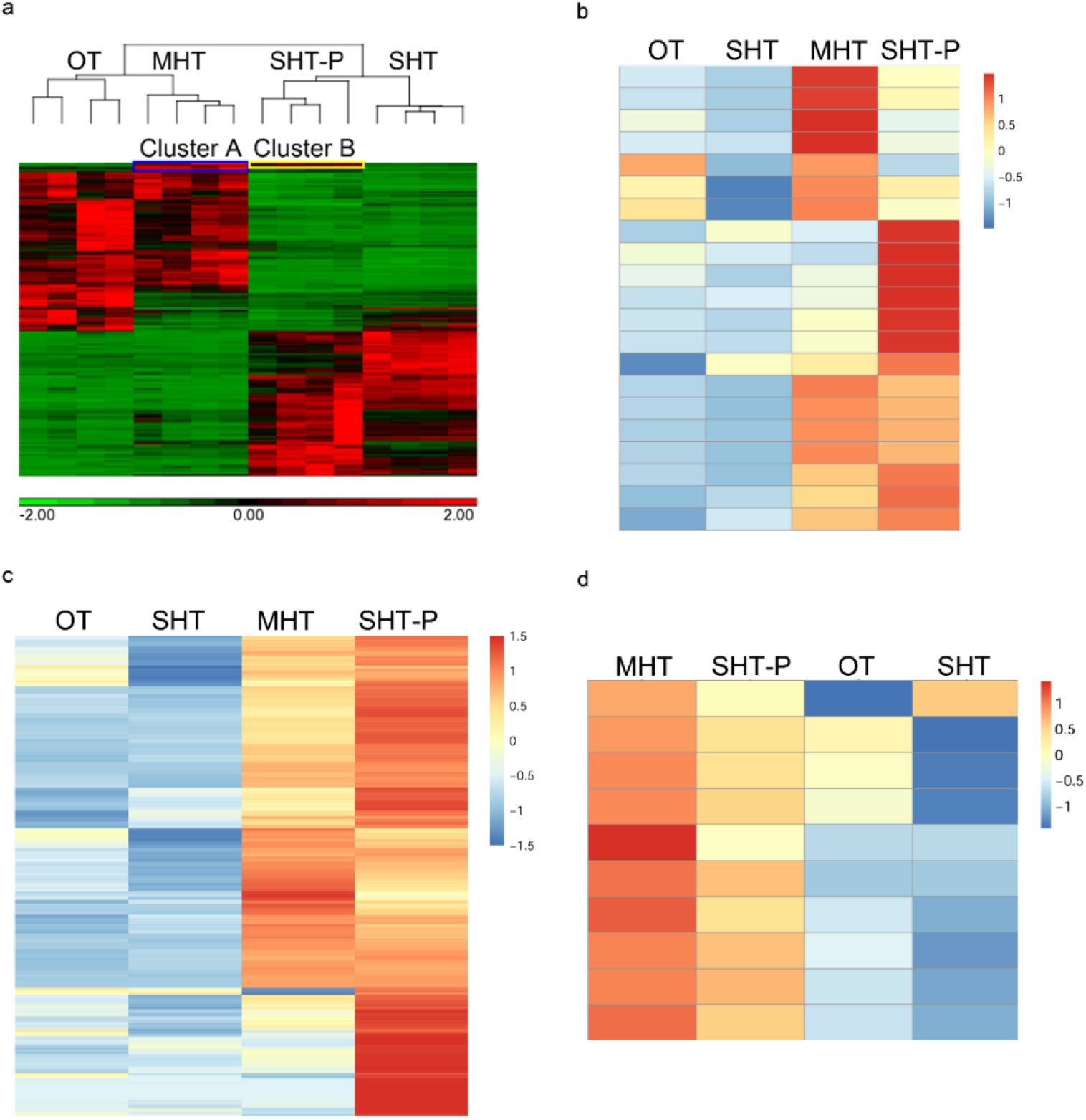
Priming-inducing genes and proteins. **a**. Expression heatmap of the 1,329 DEGs (pFDR < 0.5, lFCl ≥ 5) identified by RNA-seq analysis. Genes in clusters A (blue box, 39 genes) and B (yellow box, 16 genes) were specifically upregulated under priming conditions: 23 out of 39 genes in cluster A were upregulated only at MHT, and genes in cluster B (16 genes) were upregulated at MHT and SHT-P. Six and four serine-type peptidase (GO:0008236) encoding genes were found in clusters A and B, respectively. **b**. Expression heatmap of the 21 serine-type peptidase genes in the *B. cinerea* genome. The mean RPKM value of each treatment (four biological replications) was used to calculate the relative expression level (*Z*-score). The heatmap was generated using R. **c**. Expression heatmap of the 355 soluble candidate priming proteins. Two criteria were applied to select priming-associated candidates among the soluble proteins identified by proteomic analysis: (i) {log_2_(SHT-P/OT)} – {log_2_(SHT/OT)} ≥ 1, to select proteins that are more abundant at SHT-P than at SHT; (ii) log_2_(SHT-P/OT) > 1 and log_2_(SHT/OT) < 1, to select proteins that are more abundant at SHT-P but not at SHT compared to OT. **d**. Expression heatmap of the 10 serine-type peptidases identified by proteomic analysis (*q*-value < 0.05, lFCl ≥ 2). The mean log_2_LFQ value of each protein in each treatment (three biological replications) was used to calculate the relative expression level (*Z*-score) and to generate the heatmaps in **c** and **d** using R.

As several serine type peptidase-encoding genes were specifically upregulated under priming conditions, we analyzed the transcript profiles of all 21 annotated *B. cinerea* serine-type peptidase genes by heatmap analysis. All 21 genes were upregulated under MHT or SHT-P, whereas most were expressed at low levels under OT and SHT (Fig. 3b).

#### Proteomic analysis

We grew fungi as described in Table S6, extracted soluble and aggregated proteins (Fig. S4a), separated the proteins by SDS-PAGE and stained them with Coomassie brilliant blue. Significantly more protein aggregates formed under SHT and SHT-P compared to OT and MHT, but minor differences were also observed between MHT and OT (Fig. S4b). We then analyzed the soluble and aggregated proteins using liquid chromatography with tandem mass spectrometry (LC-MS/MS). We detected 4,492 proteins (Table S7), with good separation between soluble and aggregated proteins as well as the four heat treatments (Fig. S4c). The aggregated proteins under SHT-P and SHT clustered together, whereas the soluble proteins at MHT and SHT-P formed a subcluster. These results suggest that priming mainly affects the amount and nature of proteins that remain soluble under heat stress. Therefore, we subsequently analyzed only the soluble proteins.

The levels of 1,374 soluble proteins (Table S8) were significantly altered (*q*-value < 0.05, lFCl ≥ 2) under the three heat treatments compared to the control (Fig. S4d). The levels of 147 proteins were higher under SHT vs. OT, a number significantly lower than those under MHT (429 proteins) and SHT-P (584 proteins). We set two criteria for identifying priming-associated candidates among soluble proteins: (i) {log_2_ (SHT-P/OT)} – {log2 (SHT/OT)} ≥ 1 for proteins more abundant at SHT-P than at SHT; and (ii) log_2_(SHT-P/OT) > 1 and log_2_(SHT/OT) < 1 for proteins more abundant at SHT-P but not SHT compared to OT. This analysis yielded 355 proteins (Fig. 3c and Table S9), which we considered to be candidate regulators (MHT-specific) or executors (MHT and/or SHT-P) of the priming response.

GO enrichment analysis of the 355 priming candidates (Table S10) highlighted several major categories, including transferases (GO:0000030 and GO:0016758), transporters (GO:0005347 and GO:1901505), and serine-type peptidases (GO:0008236). Serine-type peptidases (STPs) were of particular interest since this group was also revealed as priming-related by RNA-seq (Fig. 3b). Except for *bcin_16g02790*, all 9 STPs identified among the 1,374 soluble proteins were more abundant at MHT and SHT-P, but not at SHT vs. OT (Fig. 3d). The six serine-type peptidases identified by RNA-seq were included in this set.

Collectively, the transcriptomics and proteomics analyses revealed subsets of priming-related GO categories that only partly overlapped. We also noticed a significant difference between changes in genes and protein levels during priming: compared with control, the number of DGEs was lowest in MHT (Fig. S3d and S3e)., whereas the changes in soluble proteins were lower at SHT (Fig. S4d).

### Changes in soluble protein abundance are only partly dependent on gene expression levels

To extract more information from our data, we compared the gene expression and protein abundance data. For each treatment, we selected all shared genes and soluble proteins with pFDR < 0.05 and evaluated the correlation between relative gene expression level and protein abundance. The correlations were similar under MHT (*R* = 0.631) (Fig. 4a) and SHT (*R* = 0.594) (Fig. 4b) and somewhat smaller under SHT-P (*R* = 0.477) (Fig. 4c). We also noticed a relatively high number of opposite correlations at SHT-P, namely increased protein and reduced transcript abundance. To explore the nature of these proteins, for each treatment, we selected all genes with log_2_(treatment/OT) < 0 and all proteins with log_2_(treatment/OT) ≥ 1 and calculated their proportions. Under SHT-P, 11.56% of all gene–protein pairs had negative correlations (Fig. 4d), a much larger proportion than under MHT (4.62%) and SHT (1.7%). These results suggest that the levels of many soluble proteins are regulated in a different manner under SHT-P vs. SHT.

**Fig 4.**
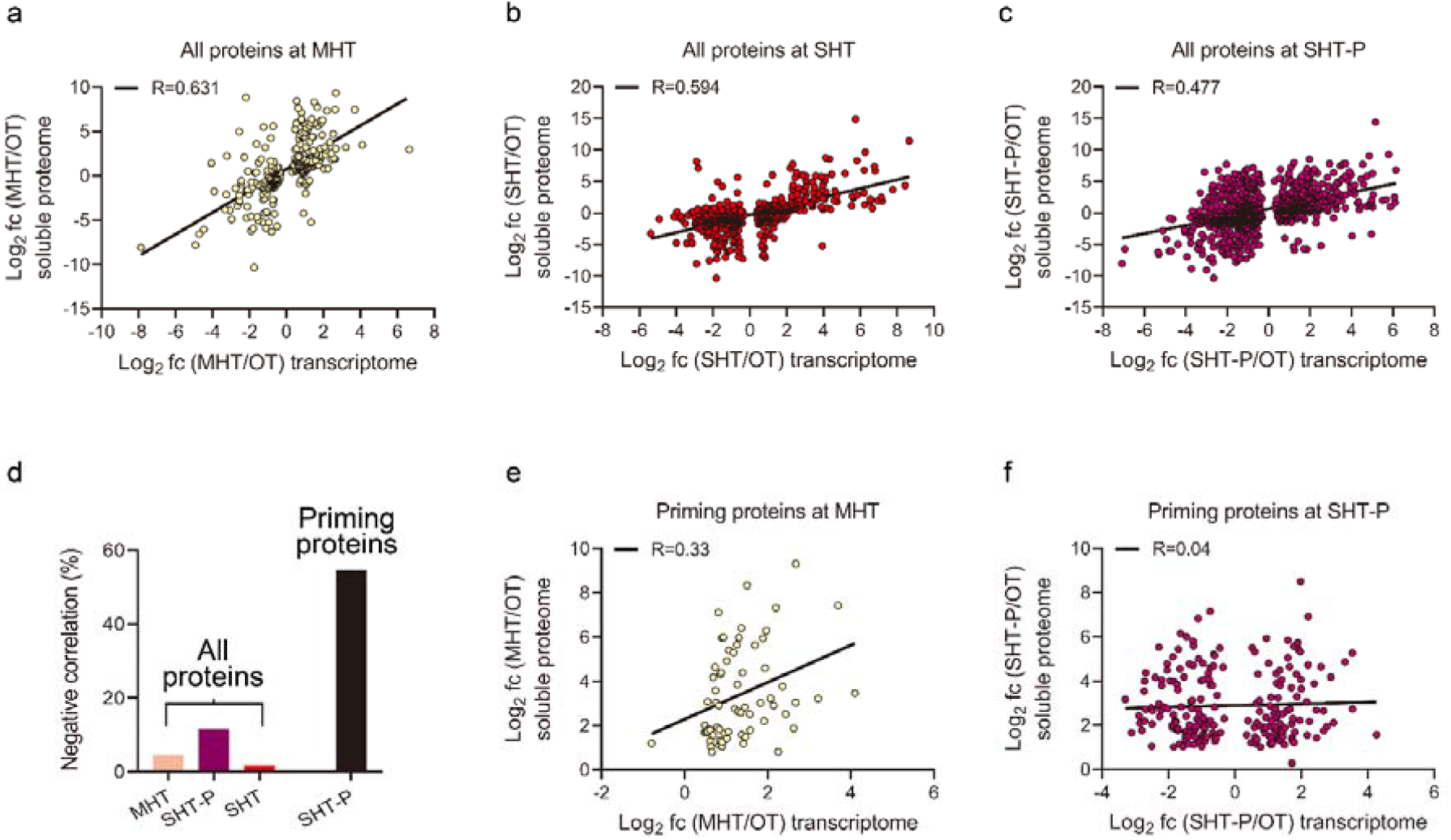
Correlations between changes in gene expression and protein levels. **a-c**. Correlation between gene expression (pFDR < 0.05) and soluble protein levels (pFDR < 0.05) at MHT, SHT and SHT-P. Soluble proteome log_2_(FC) values were scattered against log_2_(FC) values of the corresponding genes in each sample. **d**. Proportions of proteins with negative correlations of abundance vs. transcript levels. The percentage of soluble proteins with increased abundance but reduced transcript levels among the entire set of all shared genes/proteins. **e-f**. Correlations between changes in the levels of the 355 soluble candidate priming proteins and the expression levels of the corresponding genes at MHT (**e**) and SHT-P (**f**). Soluble proteome log_2_(FC) values were scattered against log_2_(FC) of the corresponding genes in each sample. Only genes with pFDR < 0.05 were included.

We performed a similar analysis of the 355 priming candidate proteins, using only the data from MHT and SHT-P, since only 20 gene–protein pairs were found at SHT. At MHT, the correlation between transcript levels and protein abundance was positive, but it dropped from 0.631 for the entire set to 0.33 (Fig. 4e). Unexpectedly, at SHT-P, the correlation between shared genes and proteins dropped from 0.477 for the entire set to nearly zero (Fig. 4f, *R* = 0.04) for the 355 priming-specific candidate proteins. In 54.6% (Fig. 4d) of the cases, transcript levels decreased while protein abundance increased. The high proportion of negative correlations between gene expression and soluble protein abundance further suggests that the abundance of soluble priming-related proteins is controlled by mechanisms other than gene expression. STPs showed the strongest positive correlation between gene expression and soluble protein levels, and similar candidates were identified by transcriptomic and proteomic analyses (Fig. 3b and 3d). We therefore investigated whether these shared STPs function in the priming response.

### Functional analysis of priming-induced serine-type peptidases

To determine whether STPs are required for heat priming, we tested the effects on priming of PMSF, which specifically inhibits STPs (32, 33). We produced GTs at 29°C, replaced the medium with fresh medium containing 0.5 or 2 mM PMSF, transferred the samples to 37°C for 4 h and stained them with PI. Treatment with 2 mM PMSF resulted in 85% cell death vs. no cell death in untreated control GTs and <5% cell death in samples treated with 0.5mM PMSF or a 2× concentration of a general protease inhibitor cocktail (Fig. 5a and S5a). Treatment with 2 mM PMSF had almost no effect on GTs that were kept at 29°C, indicating that the drug suppressed the priming response but did not kill the fungus.

**Fig 5.**
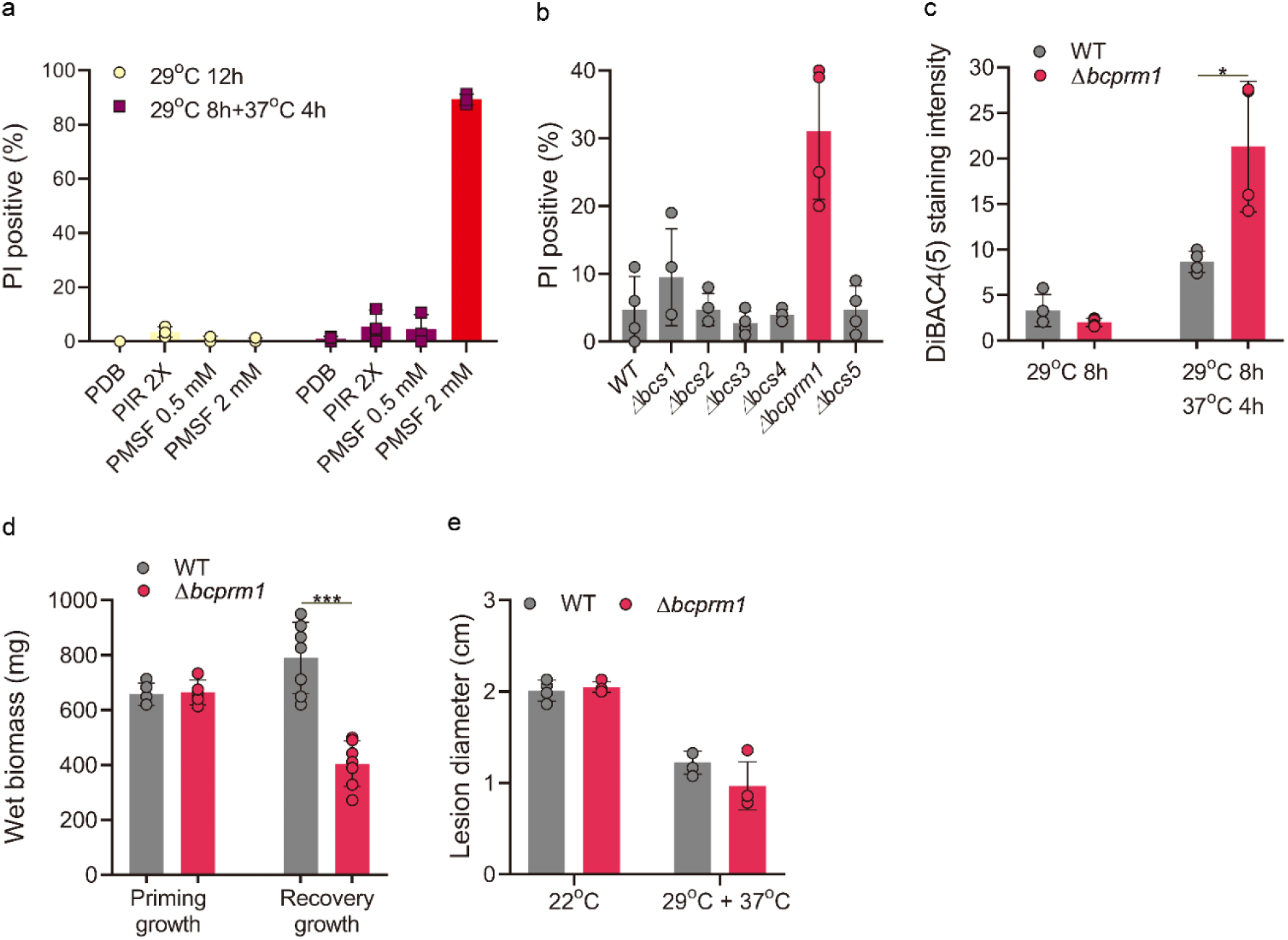
The roles of serine-type peptidases (STPs) in priming. **a**. Effects of protease inhibitors on priming. Spores were incubated in PDB at 29°C for 8 h, transferred to fresh medium with the indicated concentrations of protease inhibitors, incubated at 37°C for 4 h and stained with PI. **b-e**. Analysis of *STP* deletion mutants. **b**. Effects on priming of single deletion mutants in each of six STPs identified by RNA-seq and proteomics analyses. GTs were produced at 29°C, transferred to 37°C for 6 h and stained with PI, and the percentage of PI-positive (dead) GTs was determined. **c-e**. Analysis of the *bcprm1* (*bcin_08g02390*) deletion strain. **c**. Measurement of cell membrane potential. GTs were produced at 29°C, transferred to 37°C for 4 h and stained with DiBAC4(5). **d**. Measurement of biomass. Initial fungal biomass was produced in malt medium at 22°C and transferred to 29°C for 12 h, the wet biomass was determined (priming growth), and the biomass was incubated at 37°C for 12 h. The cultures were then transferred to 22°C for 24 h, the mycelia were collected and biomass weight was determined (recovery growth). **e**. Infection assay. Inoculated plants were incubated for 8 h at 29°C, transferred to 37°C and incubated for 4 h, and finally incubated at 22°C for 3 d and the lesion diameter measured. Graphs represent four (**a**-**c**,**e**) or at least five (**d**) biological replications with overlaid individual data points. Values are presented as the mean of replicates ± s.d. Statistical differences were determined according to unpaired two-tailed Student’ t-test (\**P* < 0.05; \*\*\**P* < 0.001).

To test the possible roles of the six identified STPs in priming, we generated deletion strains for each of the six genes and tested their priming responses. All deletion strains had normal colony morphology, hyphal growth and germination rates at 22°C and 29°C (not shown). To evaluate priming responses, we produced GTs at 29°C, transferred them to 37°C and stained them with PI. Compared to the wild type, strain Δ*bcin_08g02390* showed markedly increased cell death, whereas the five other deletion strains showed normal levels of cell death (Fig. 5b and S5b). Similarly, the membrane potential of strain Δ*bcin_08g02390* was compromised following transfer from 29°C to 37°C (Fig. 5c and S5c).

Four of the STPs belong to peptidase group S53 (*BCIN_06g00620, BCIN_06g00330, BCIN_15g04670* and *BCIN_15g03150*). To determine whether the lack of a priming phenotype in a single deletion strain of these genes resulted from functional redundancy, we generated the strain Δ4stp, with deletions of all four S53 genes. Similar to the single deletions trains, the Δ4stp strain did not show developmental defects and had a normal priming response (not shown). Hence, the role of these four S53 genes in priming remains unverified.

Because the deletion of *bcin_08g02390* compromised the priming response of the fungus, we named this gene *bcprm1* (*B. cinerea* priming 1). To further examine the physiological relevance of the priming defects, we examined the effect of priming on biomass production of the Δ*bcprm1* strain. We produced uniform wild-type and Δ*bcprm1* cultures in malt medium at 22°C, transferred them to 29°C for 12 h to induce priming, and measured the biomass. We then incubated cultures for 12 h at 37°C followed by 22°C for 24 h, and measured their biomass once again (Fig. S5d). There were no differences in biomass between the wild-type and Δ*bcprm1* strains after incubation at 29°C. However, the recovery of Δ*bcprm1* after incubation at 37°C was compromised, as it produced ∼50% less biomass than the wild type (Fig. 5d and S5e). To examine the effect of the deletion of *bcprm1* on pathogenicity, we inoculated the leaves of *P. vulgaris* plants with a spore suspension and incubated them at 29°C for 8 h, then at 37°C for 4 h and then at 22°C for 3 d. Compared to the wild type, plants inoculated with the mutant strain developed less severe symptoms (Fig. 5e and S5f). Collectively, these results indicate that the STP BcPrm1 is required for a full-scale priming response and contributes to fungal survival under SHTs.

## Discussion

Understanding the mechanisms of heat adaptation in fungi is essential for risk evaluation in preparation for temperature-driven disease outbreaks (34, 35) and for the rational design of temperature-driven disease control methods. To help achieve these goals, we studied the mechanisms of heat adaptation in *B. cinerea*, a cosmopolitan, devastating plant pathogen (36, 37).

We found that heat stress adaptation via priming is a powerful mechanism that enables *B. cinerea* to cope with potentially lethal temperature conditions. Comparative analysis revealed poor correlations between changes in gene expression and protein abundance under priming conditions, which were most prominent within a set of 355 priming-related soluble proteins (Fig. 3c). Muhlhofer et al. (2019) (2) reported similar results in yeast and proposed that the massive upregulation of gene expression under moderate heat stress is required to counterbalance increased protein turnover and to maintain metabolism under temperature stress. Accordingly, we propose that the main function of priming is to maintain protein solubility under SHTs by activating the compensation system, which stimulates gene expression and protein synthesis (Fig. 6). This assumption is supported by the relatively good correlation between upregulated genes and proteins under MHT (Fig. 4a). Under MHT, cellular damage is initially low, and high protein levels lead to excess cellular production and hence accelerated GT growth (Fig. S1c and S1d), as also demonstrated in yeast (2). During longer stress periods, damage slowly accumulates, cellular programs deviate from their optimal functions and mycelial growth decreases over time (Fig. S1b). The high levels of priming-induced soluble proteins might serve as a buffer that mitigates the detrimental effects of exposure to SHTs. Apart from their buffering capacity, some proteins, such as the priming-induced STPs, likely have more specific roles.

**Fig 6.**
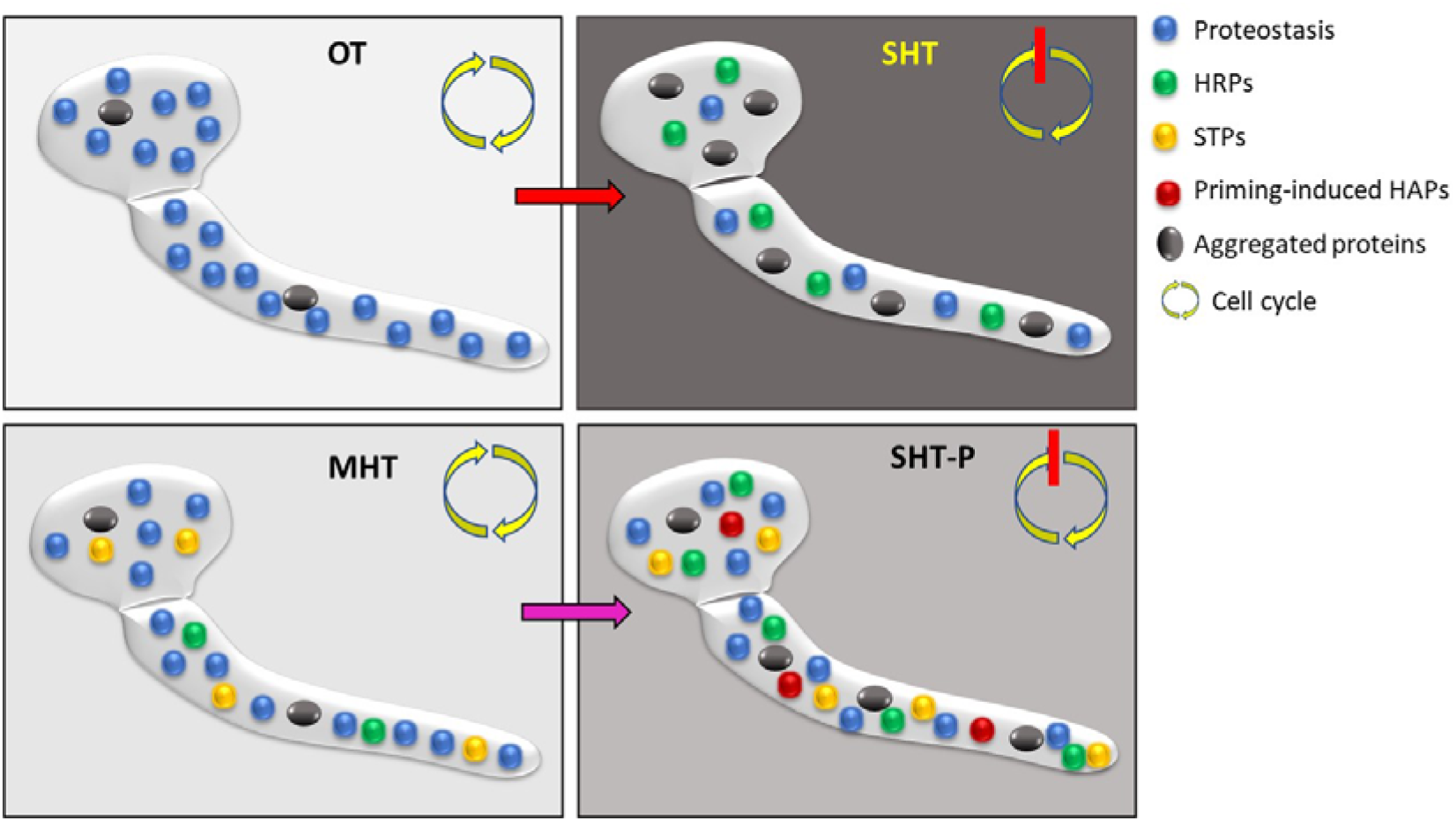
Proposed model of heat priming. Under optimal temperature (OT), the processes of protein production, aggregation and recycling are balanced and proteostasis is maintained. Moderately high temperatures (MHTs; priming temperature) induce the accumulation of priming-activating proteins, such as certain serine-type peptidases (STPs) and heat shock proteins (HSPs), protein aggregation remains low, and proteostasis and growth are maintained. A shift from OT to severely high temperature (SHT) causes an immediate arrest of the cell cycle and growth coupled with inhibited transcription and translation, accumulation of heat response proteins (HRPs), massive protein aggregation with reduced amounts of soluble proteins, loss of proteostasis and homeostasis, and rapid cell death. A shift from MHT to SHT (SHT-P) also causes cell cycle arrest, reduced protein synthesis and the accumulation of protein aggregates. However, this is accompanied by the rapid accumulation of heat adaptation proteins (HAPs) such as STPs, HSPs and trehalose biosynthesis enzymes. Therefore, a relatively high level of soluble proteins is maintained, proteotoxicity is reduced and the accumulation of cellular damage is slowed, thereby extending the survival time of the fungus under SHT.

Several lines of evidence, including transcriptomics, proteomics, pharmacological and mutagenesis data, link STPs to the *B. cinerea* priming response, suggesting that STPs are an important group of heat adaptation proteins. STPs are highly abundant in all organisms (38) and are involved in proteostasis, thereby contributing to cell fitness and survival (9, 39). In *Arabidopsis thaliana*, the extracellular subtilase SBT3.3 is required for the activation of immune priming, which is mediated by a chromatin-remodeling- and salicylic-acid-dependent mechanism (40). Specific serine peptidases, such as the HTRA (41, 42), Clp (43, 44) and Lon1 (8, 9) proteases, are associated with heat adaptation, possibly through stabilization of specific proteins or removal of stress-induced protein aggregates. The upregulation of the six STP genes during both MHT and SHT-P treatment suggests that STPs might serve as both sensors for priming induction and executors of the priming response, possibly facilitating protein solubility and disaggregation.

Our results highlight the importance of priming in mediating the heat adaptation of fungi and advance our understanding of how psychrophilic fungi manage to survive in hot environments. We demonstrated the potential application of this knowledge by subverting the priming response through a protocol involving a series of temperature shifts, which eliminated the fungus and prevented disease development. A deeper understanding of specific elements that regulate priming, such as the mode of action of STPs and a system-level understanding of the priming response, will improve our ability to target this mechanism for disease management.

## Materials and Methods

### Fungal cultures

*B. cinerea* strain B05.10 and derived transgenic strains were routinely cultured on potato dextrose agar (PDA, Acumedia) or suspended in potato dextrose broth (PDB) at 22°C under continuous fluorescent light supplemented with near UV light. Additional media that were used for specific experiments included malt medium (5 g glucose, 15 g malt extract, 1 g peptone, 1 g casamino acids) and GB5-Glc medium (Gamborg B5 with vitamins and 2% glucose, Duchefa Biochemie).

### Germination, colony growth, and germ tube elongation

#### Germination

Fungi were cultured on PDA for 7 d. Spores were collected by washing with PDB and filtered through two layers of Miracloth, and the spore density was adjusted to 5 × 10^5^/ml. A 20-μl droplet of spore suspension was placed on a coverslip and incubated under continuous light at the specified temperature and time. After incubation, the slides were visualized under a light microscope and germination rates were scored. Each experiment was repeated at least four times, with >200 randomly selected spores each time.

#### Colony growth

Cultures were initiated from 4-mm plugs that were cut from the edge of a 2-day-old colony. The plugs were placed in the center of a Petri dish containing PDA and incubated at the indicated temperatures under continuous light. Colony diameter was recorded after 3 d, the diameter of the initial inoculation plug (4 mm) was subtracted, and radial growth was calculated. Each experiment was performed with three replications (3 plates per treatment) and was repeated three times.

#### Germ tube elongation

A 20-μl droplet of spore suspension (5 × 10^4^/ml) was placed on a coverslip and incubated under the indicated conditions. After incubation, the slides were visualized under a light microscope and GT length was recorded. Each experiment was repeated three times, with more than 120 randomly selected GTs each time.

#### Germination after heat stress

Fungi were cultured on PDA for 7 d to allow mycelia and spores to develop. The plates were transferred to 37°C or 42°C for the indicated time. After heat treatment, spores were collected into PDB, and spore density was adjusted to 200 spores/ml. To determine germination rates, a 5-μl spore suspension was placed on a small (1 cm^2^) PDA cube and incubated at 22°C for 72 h. The number of germinated and un-germinated spores was scored, and germination rate was calculated. Each experiment was performed at least three times with >200 spores per treatment.

### Priming-related assays

#### Cell death

Spores (2 × 10^5^/ml PBD) were placed on a coverslip, incubated at 22°C or 29°C for 12 h under continuous light and transferred to 37°C. Following incubation at 37°C, the GTs were stained with 10 µg/ml PI for 15 min. Samples were visualized under a Zeiss Axio imager M1 fluorescence microscope using a rhodamine filter. The number of dead cells (PI-positive) was counted, and the proportion of dead cells was calculated. Each experiment was repeated at least three times with more than 120 randomly selected GTs each time.

#### Membrane potential

GTs were suspended in PBS containing 100 µM DiBAC4(5) (Interchim), incubated for 15 min at room temperature and visualized under a fluorescence microscope using a rhodamine filter. Images were captured using Zeiss AxioCam MRm camera. The fluorescence intensity of each germ tube was quantified using ImageJ, and the mean signal intensity of each treatment was calculated. The experiment was repeated four times with at least 50 randomly selected GTs each time.

#### Recovery of colony growth after heat shock

Colonies were initiated by placing a 5-μl droplet of spore suspension (5 × 10^5^/ml) in the center of a Petri dish containing PDA. The plates were incubated under continuous light at 22°C or 29°C for 12 h and then at 37°C for 6 or 9 h. After heat treatment, the cultures were incubated at 22°C for 3 d and the colony diameter measured. Each experiment was repeated six times with five replications per treatment.

#### High/low temperature cycles

Spores (2 × 10^5^/ml) were incubated in liquid GB5-Glc on a coverslip at 22°C for 6 h, transferred to 42°C for 2 h (heat treatment) and incubated at 22°C for 22 h (recovery). The cycle was repeated three times, and the levels of cell death were determined after each cycle by PI staining. The experiments were repeated at least three times with >300 randomly selected GTs each time.

#### Infection assays

Pathogenicity assays were performed using the first two leaves of 8-day-old French bean (*Phaseolus vulgaris* L. genotype N9059) plants as described previously (45). The leaves were inoculated with 7.5-μl droplets of a spore suspension containing 2 × 10^5^ spores/ml in GB5-Glc. The plants were incubated in closed boxes at 100% relative humidity, and disease levels were estimated visually by scoring lesion diameter at the indicated time post inoculation.

For priming experiments, inoculated plants were incubated at 22°C or 29°C for 16 h and transferred to 37°C for 4 or 8 h. Following the heat treatment, the plants were incubated at 22°C for 3 d and the lesion diameter measured. Each experiment was repeated at least four times with three plants (two leaves per plant) per treatment.

To evaluate the effects of high/low temperature cycles on pathogenicity, inoculated plants were first incubated at 22°C for 6 h to allow spores to geminate and produce GTs, then transferred to 42°C for 2 h, followed by 22°C for 22 h. This cycle was repeated three times. The plants were then incubated at 22°C for three more days. The experiments were repeated five times with three plants (two leaves per plant) per treatment.

### DNA and RNA extraction and analysis

Genomic DNA extraction was performed using Extract-N-Amp(tm) Tissue PCR Kits (Sigma/Aldrich). For cDNA synthesis, total RNA was treated with DNase I (Thermo Scientific) and first-strand cDNA was synthesized from 1μg of DNA-free RNA using a RevertAid First Strand cDNA Synthesis Kit (Thermo Scientific). qRT-PCR was performed with SYBR Premix Ex Taq II (Takara, Dalian, China) using a StepOne (Applied Biosystems) Real-time PCR instrument (Applied Biosystems). Relative fold changes of mRNA levels determined by RNA-seq were normalized to the expression of the ribosomal protein gene *bcrsm18*, which has relatively stable mRNA expression.

### RNA-seq and analysis

#### Biological materials

*B. cinerea* cultures were produced by spreading 500 μl of spore suspension (10^7^/ml) on a cellophane-covered PDA plate. Treatments (Table S1) included incubation at 22°C or 29°C for 10 h (OT and MHT, respectively), and 22°C or 29°C for 8 h followed by 37°C for 2 h (SHT and SHT-P, respectively). GTs were collected from the plates by washing with water, centrifuged, separated into two 1.5-ml Eppendorf tubes, flash-frozen in liquid nitrogen and stored at –80°C. RNA was extracted from one tube per sample, with four biological replications per treatment and a total of 16 samples. Samples were ground in liquid nitrogen with a mortar and pestle, and RNA was extracted with TRIzol reagent (Sigma-Aldrich) according to the manufacturer’s instructions.

#### RNA quality, library preparation, and RNA-seq

cDNA libraries were prepared using a NEBNext Ultra II RNA Library Prep Kit (NEB). RNA and cDNA quality and quantity were evaluated by performing a Qubit assay with the TapeStation 4200 system (Agilent Technologies). Sequencing libraries were constructed with barcodes using NEBNext multiplex oligos for Illumina (NEB). Pooled libraries of 16 samples were sequenced at the Faculty of Life Sciences, Tel Aviv University, Israel in one lane of the Illumina NextSeq 500 platform using the 75-bp single-end RNA-seq protocol.

Raw reads were uploaded to the Partek Flow commercial tool (build version 9.0.20.0804), and 3’-end low-quality bases (<Phred 20) were trimmed. An average of more than 24 million reads per sample was obtained, with high quality (Phred score > 34). The number of reads after QC and trimming ranged from 19,604,233 to 29,043,828 (Table S2). The clean reads were mapped to the *B. cinerea* reference genome (assembly ASM14353v4, https://www.ncbi.nlm.nih.gov/genome/494) using STAR-2.7.3a. Over 97% of the reads in each sample were mapped to the *B. cinerea* genome (Table S2), and 11,219 of the 11,700 annotated *B. cinerea* strain B05.10 genes were identified. Principal component analysis (PCA) separated the samples into four distinct clusters based on treatment (Fig. S3a). RT-PCR analysis of eight genes validated the RNA-seq results, with highly similar expression patterns, confirming the accuracy of the RNA-seq data (Fig. S3b and S3c). Quantification and normalization of gene expression were performed using the Partek E/M algorithm and DESeq2, respectively. Gene expression values were calculated as fragments per kilobase per million (FRKM). Principal coordinate analysis and heatmap analysis were performed using the Partek Flow tool or R software. Identification of DEGs was performed with the DESeq2 package with cutoffs of pFDR < 0.05 and |fold change| (FC) ≥ 2 or ≥ 5. GO enrichment analysis of the DEGs was performed using the FungiDB platform (https://fungidb.org/fungidb/app) (46).

### Isolation of soluble and aggregated proteins

To isolate soluble and aggregated proteins, fungal material was produced as described for the RNA-seq experiment, except that the initial incubation at 22°C or 29°C was performed for 14 h instead 8 h to allow the accumulation of sufficient biomass. Soluble and aggregated proteins were extracted from the samples according to Koplin and Brennan (47, 48) with some modifications. At the end of the heat treatments, the fungal biomass was collected (0.8–1.6 g), ground to a fine powder in liquid nitrogen and suspended in 1.2 ml of freshly prepared protein extraction buffer (20 mM Na-phosphate, pH 6.8, 10 mM DTT, 1 mM EDTA, 0.1% Tween, 1 mM PMSF, 1× Mini Protease Inhibitor Cocktail, Roche). The samples were centrifuged at 3000*g* for 6 min at 4°C using a tabletop centrifuge, and each supernatant was transferred to a new 1.5-ml Eppendorf tube. The samples were centrifuged at 5,000*g* for 5 min at 4°C to precipitate remnants of cellular debris, and the supernatant was transferred to a new tube. A sample of the protein extract was measured in a spectrophotometer, and the amount of total protein in each sample was calculated using a 1 mg/ml standard sample of bovine serum albumin (Sigma-Aldrich). To separate the soluble and aggregated proteins, extracted proteins were centrifuged at 16,000*g* for 20 min at 4°C, and the supernatant was carefully removed, transferred to a new tube and stored at –80°C as the soluble protein fraction. The pellet was washed twice with 300 μl sodium phosphate buffer (20 mM Na-phosphate pH 6.8, 1 mM PMSF, 1× Mini Protease Inhibitor Cocktail, Roche) with centrifugation at 16,000*g* for 20 min between washes. The supernatant was removed and the clean pellet stored at –80°C as the protein aggregate fraction.

### SDS-PAGE and proteomic analyses

For soluble proteins, samples were mixed with protein loading buffer and boiled for 5 min. For protein aggregates, the protein pellet was dissolved in 20 μl of 8 M urea, mixed with protein loading buffer and boiled for 5 min. The amounts of proteins were normalized to represent an equal amount of total proteins in all samples. The protein samples were separated by SDS-PAGE and stained with Coomassie brilliant blue.

Proteomics analysis was performed at the Smoler Proteomics Center, the Technion, Israel as previously described (49). Samples were incubated at 60°C for 30 min to reduce the volume, mixed with 35 mM iodoacetamide in 400 mM ammonium bicarbonate and incubated at room temperature in the dark for 30 min. The samples were then mixed with 2 M urea, 80 mM ammonium bicarbonate solution and digested with modified trypsin (Promega) by overnight incubation at a 1:50 enzyme-to-substrate ratio at 37°C. The resulting tryptic peptides were desalted using C_18_ tips (Harvard), dried and resuspended in 0.1% formic acid. The samples were analyzed by LC-MS/MS using a Q Exactive Plus mass spectrometer (Thermo) fitted with a capillary high-performance liquid chromatograph (HPLC; easy nLC 1000; Thermo). The peptides were loaded onto a homemade capillary column (25-cm, 75-μm internal diameter) packed with Reprosil C_18_ Aqua (Dr. Maisch GmbH, Ammerbuch, Germany) in solvent A (0.1% formic acid in water). The peptide mixture was resolved with a linear gradient (5%–28%) of solvent B (95% acetonitrile with 0.1% formic acid) for 105 min, followed by a 15-min gradient of 28%–95% and 15 min at 95% acetonitrile with 0.1% formic acid in water at a flow rate of 0.15 μl/min. Mass spectrometry was performed in positive mode (*m*/*z* 350 to 1,800; resolution 70,000) using a repetitively full MS scan, followed by collision-induced dissociation (high-energy collision dissociation [HCD], at a normalized collision energy of 35) of the 10 most dominant ions (>1 charges) selected from the first MS scan. The AGC settings were 3 × 10^6^ for the full MS scan and 1 × 10^5^ for the MS/MS scans. The intensity threshold for triggering MS/MS analysis was 1 × 10^4^. A dynamic exclusion list was enabled with an exclusion duration of 20 s.

The mass spectrometry data from all three replications were analyzed using MaxQuant software v.1.5.2.8. (50) for peak identification and quantitation using the Andromeda search engine, which searches for tryptic peptides against the B05.10 UniProt database (51), with a mass tolerance of 20 ppm for both the precursor masses and fragment ions. Oxidation on methionine and protein N-terminus acetylation were accepted as variable modifications, and carbamidomethyl on cysteine was accepted as a static modification, as the percentage of carbamylation was low. Minimal peptide length was set to 6 amino acids, and a maximum of two missed cleavages were allowed. Peptide- and protein-level false discovery rates (FDRs) were filtered to 1% using the target decoy strategy. Protein tables were filtered to eliminate identifications from the reverse database and common contaminants and single peptide identifications. The data were quantified by SILAC analysis using the same software. H/L ratios for all peptides belonging to a particular protein species were pooled, providing a ratio for each protein.

### Protease inhibitor assay

A 20-μl droplet of spore suspension (10^5^/ml) was placed on a coverslip and incubated at 29°C for 8 h. The suspension medium was carefully removed and replaced with fresh PDB supplemented with 2× protease inhibitor (Mini Protease Inhibitor Cocktail, Roche), 0.5 mM or 2 mM PMSF (Sigma-Aldrich), or an equal volume of PDB (control treatment). The samples were incubated at 29°C or 37°C for 4 h, the GTs were stained with PI, and the death rate was calculated. Each experiment was repeated at least three times with more than 150 randomly selected GTs per sample.

### Generation of serine protease deletion strains

Deletion strains were generated using a marker-free CRISPR-Cas9 genome editing method (52) with two sgRNAs per gene. All CRISPR-Cas9 reagents were purchased from Integrated DNA Technology (IDT). Selection and design of sgRNAs were conducted with a sgRNA design web platform (http://grna.ctegd.uga.edu/). To assemble a single sgRNA duplex (33µM), 0.2 nm each of Alt-R CRISPR-Cas9 crRNA (2µl) and Alt-R CRISPR-Cas9 tracrRNA (2µl) were mixed with Nuclease-Free Duplex Buffer (2 µl). The mixture was incubated for 5 min at 95°C and allowed to cool at room temperature. To assemble a Cas9/sgRNA ribonucleoprotein (RNP) complex, 3 μg Cas9 (3 μl), 33 µM sgRNA duplex (2 µl) and 5 µl of Cas9 working buffer were mixed and incubated at 37°C for 30 min.

Transformation of *B. cinerea* was performed according to Leisen et al. (53). Briefly, protoplasts were incubated with 2 µg pTEL-Fen telomeric plasmid (52) and two RNP complexes per gene, each containing 3 µg Cas9 and 1 µg sgRNA. The protoplasts were mixed with 50 ml of liquified SH agar medium supplemented with 30 mg/l fenhexamid (Fen; Teldor), and the mixture was dispensed into 90-mm Petri dishes. The plates were incubated at 22°C for 3 d, and Fen-resistant colonies were collected. The colonies were transferred to PDA plates without selection to allow rapid growth and the loss of pTEL-Fen selection. Hyphal tips from fast-growing isolates were transferred to fresh PDA plates, allowed to grow for 4 d and subjected to DNA extraction and PCR analysis to verify the deletion. A single round of single spore isolation was performed, followed by DNA extraction and PCR analysis to verify that the strains were homokaryotic for the deletion. To generate strains with deletions of multiple genes, purified strains were selected and transformed with RNP complexes for additional genes. For each strain, three to four individual isolates were selected and subjected to initial phenotyping by examining growth morphology, sporulation, spore germination, and the priming responses of GTs.

### Biomass production

Liquid cultures were initiated by inoculating 50 ml malt medium with 10^6^ spores/ml. The cultures were incubated for 12 h at 22°C with shaking at 150 rpm under continuous light and transferred to 29°C. After 12 h incubation with shaking at 29°C, two samples of 10 ml each were removed, placed in 15-ml tubes and centrifuged at 4000*g* for 15min. The supernatant was discarded and the fresh weight was measured. The tubes with pellets were incubated in an oven at 55°C for 24 h and the dry weight measured. After removing 20-ml aliquots, the remaining cultures were transferred to 37°C and incubated 12 h, followed by 22°C for 24 h. A 20-ml aliquot was removed from each sample, and the fresh and dry weights were measured. Each experiment was repeated three times with two replications per treatment.

### Statistical analysis

The statistical significance between means of treatments was evaluated by Student’s *t*-test (two-tailed *t*-test). In all graphs, results represent the mean values of at least three independent experiments, each with at least three replications per treatment. Details of statistical analyses are presented in the figure legends and figures were generated using Prism software (GraphPad 8.0).

## Author contributions

Conceptualization: A.S. and M.Z. Methodology: A.S., M.H., and M.Z. Investigation: M.Z., N.T., and T.L. Visualization: M.Z. and M.C. Supervision: A.S. Writing— original draft: A.S. and M.Z. Writing—review & editing: A.S. and M.Z.

## Data and materials availability

All data needed to evaluate the conclusions in the paper are present in the paper and/or the supplementary materials. RNA sequencing data have been uploaded to NCBI and will be available once granted a project accession number. The mass spectrometry proteomics data have been deposited in the ProteomeXchange Consortium via the PRIDE (54) partner repository with the dataset identifier PXD037173. Additional data are available from the corresponding authors upon request.

---

**Fig S1.**
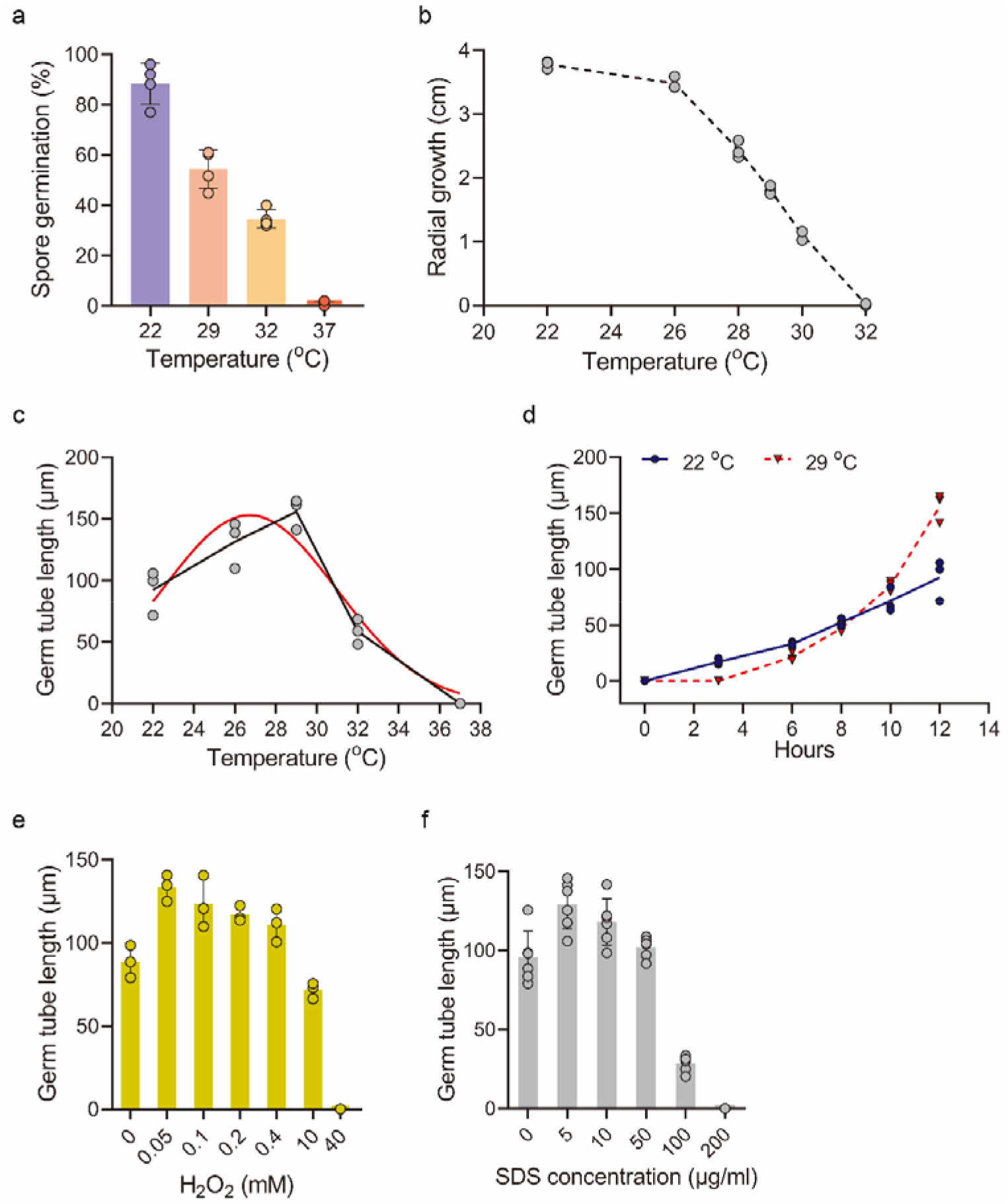
Growth responses of *B. cinerea* to different temperatures. **a**. Spore germination. Spores were incubated at the indicated temperature, and germination rates were determined after 3 h of incubation at 22°C, 4 h at 29°C, 8 h at 32°C or 12 h at 37°C. **b**. Colony growth. Colonies were initiated from mycelial plugs, and radial growth was measured after 3 d of incubation at the indicated temperatures. **c**. Germ tube growth. Spores were incubated at the indicated temperatures, and GT length was measured after 12 hours of incubation. Red line shows a Gaussian regression curve (*R*^2^ = 0.9186). **d**. Germ tube length over time. Spores were incubated at 22°C or 29°C, and GT length was recorded at different time intervals. **e**,**f**. Effects of oxidative (H_2_O_2_) and cell wall (SDS) stress on GT growth. Spores were incubated in medium with the indicated concentrations of H_2_O_2_ or SDS, and GT length was recorded after 12 h. Graphs represent four (**a**), three (**b**–**e**) and six (f) biological replications with overlaid individual data points. For **a, e** and **f**, values are presented as the mean of replicates ± s.d.

**Fig S2.**
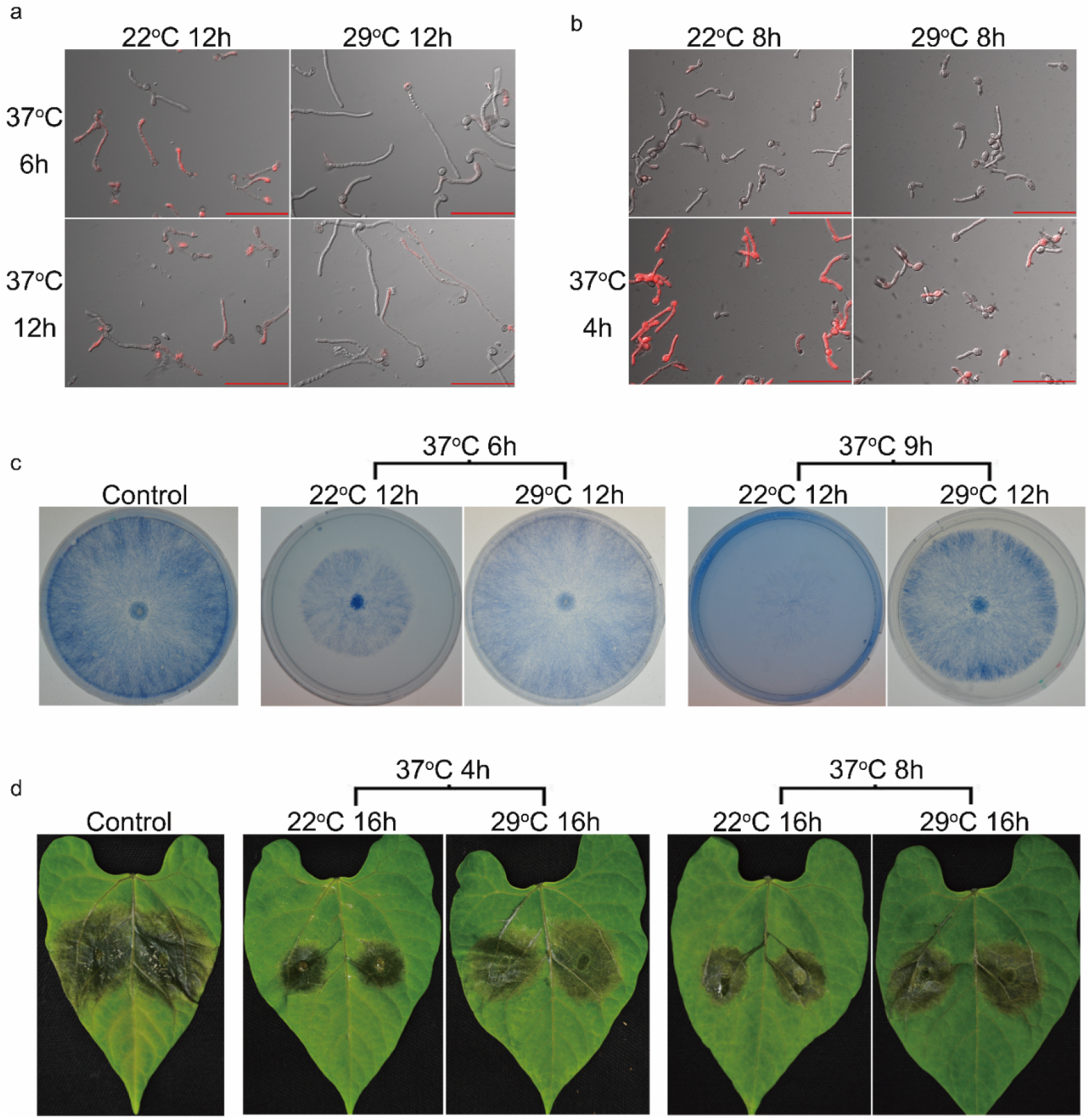
Effects of priming on fungal survival, recovery, and pathogenicity. **a**,**b**. Combined DIC and fluorescence (rhodamine filter) microscopic images of GTs after staining with PI (**a**) or DiBAC4(5) (**b**). Scale bars, 100 μm. **c**. Recovery growth of fungal cultures after exposure to 37°C with and without priming. Cultures were stained with cotton blue for better visualization of mycelia. **d**. Infection symptoms of inoculated French bean (*Phaseolus vulgaris*) leaves following exposure to 37°C with and without priming.

**Fig S3.**
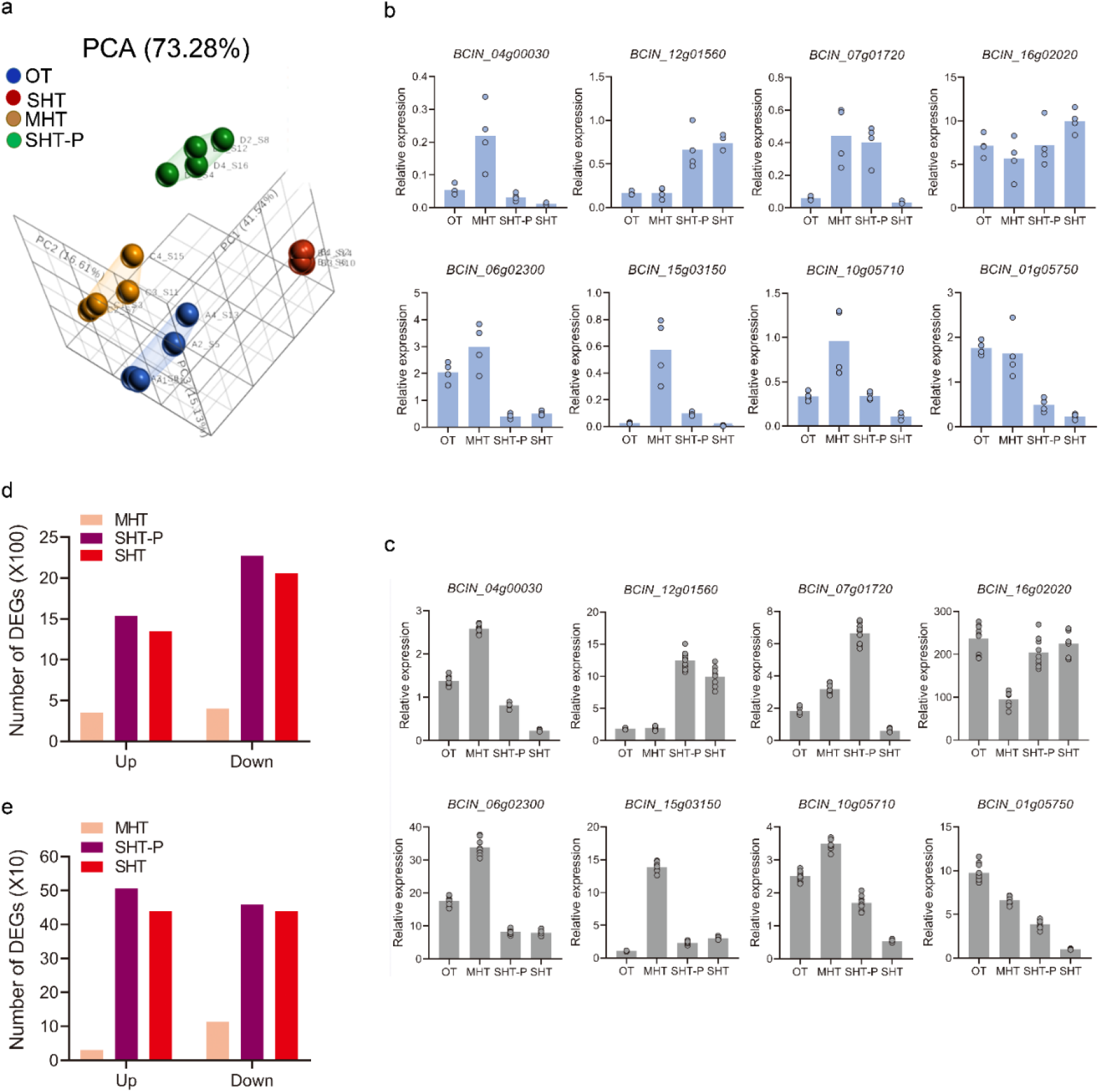
RNA-seq data. **a**. Principal component analysis (PCA) of all genes that were identified by RNA-seq. Fungi were grown under optimal temperature (OT, 22°C), moderately high temperature (MHT, 29°C), severely high temperature (SHT, 22°C + 37°C) or severely high temperature conditions with priming (SHT-P, 29°C + 37°C). Mycelia were harvested, and RNA was extracted and sequenced in one lane of the Illumina NextSeq 500 flow cell using the 75-bp single-end RNA-seq protocol. **b**. Relative expression levels of eight genes in the RNA-seq data. Relative expression was calculated using the RPKM value under each treatment (four biological replications) and normalized to the expression of the ribosomal protein gene *bcrsm18*. Graphs represent four biological replications with overlaid individual data points. **c**. Validation of the RNA-seq results by RT-PCR analysis. Relative expression was normalized to the ribosomal protein gene *bcrsm18*. Graphs represent nine overlaid individual data points from three independent biological replicates and three technical replicates. **d**. Number of DEGs (pFDR < 0.05, lFCl ≥ 2) at MHT, SHT-P and SHT compared to OT. **e**. Number of DEGs (pFDR < 0.05, lFCl ≥ 5) at MHT, SHT-P and SHT compared to OT.

**Fig S4.**
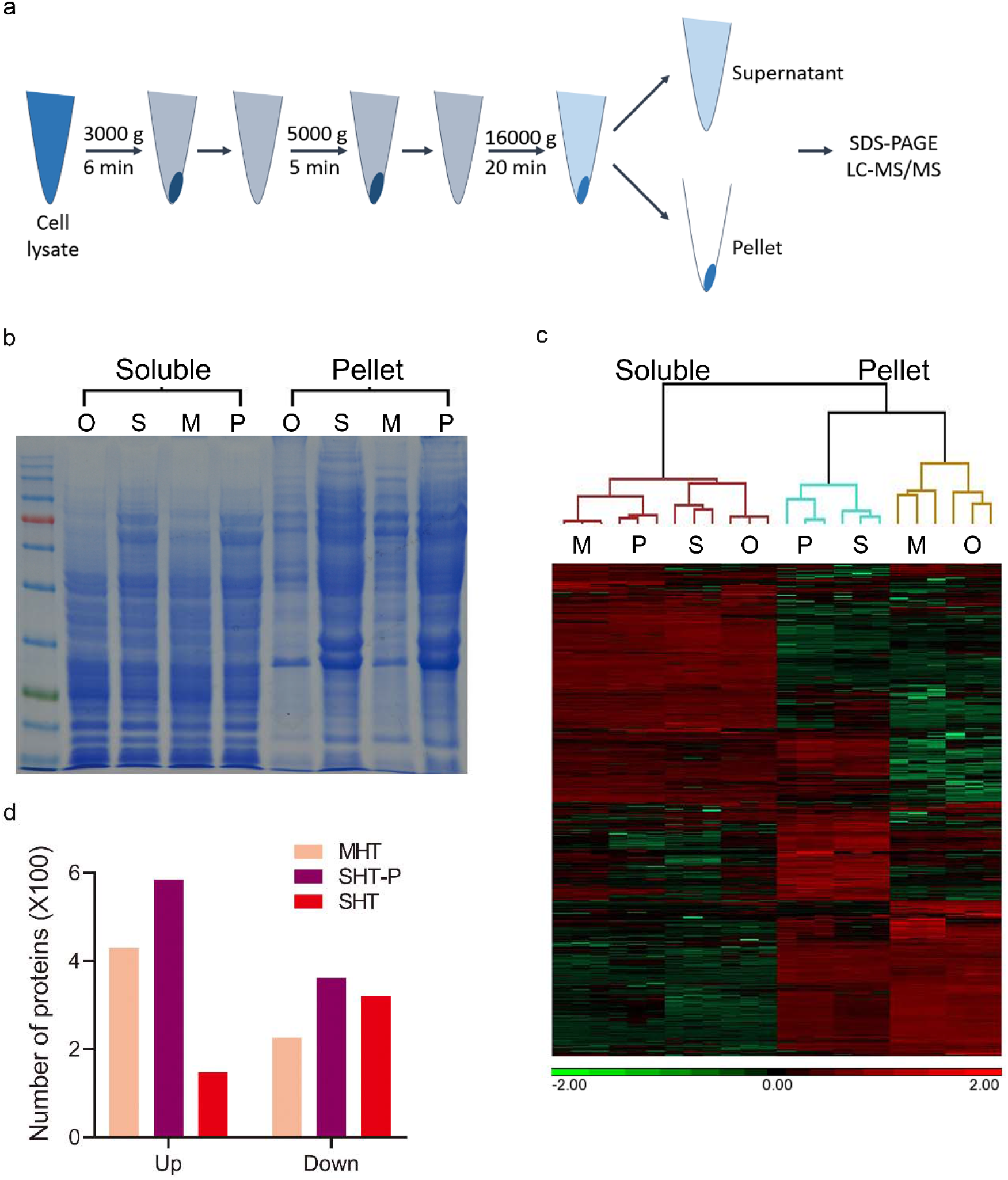
Proteomic data. **a**. Workflow for the separation of soluble and pellet proteins. **b**. SDS-PAGE analysis of Coomassie brilliant blue stained soluble (left four lanes) and pellet (right four lanes) proteins. O: OT; M: MHT; P: SHT-P; S: SHT. **c**. Expression heatmap of all the proteins identified by proteomics analysis. Soluble and pellet proteins were analyzed by liquid chromatography with tandem mass spectrometry (LC-MS/MS). A total of 4,492 proteins were detected. O: OT; M: MHT; P: SHT-P; S: SHT. **d**. Number of soluble proteins that were up-or downregulated (*q*-value < 0.05 and lFCl ≥ 2) at MHT, SHT-P and SHT compared to OT.

**Fig S5.**
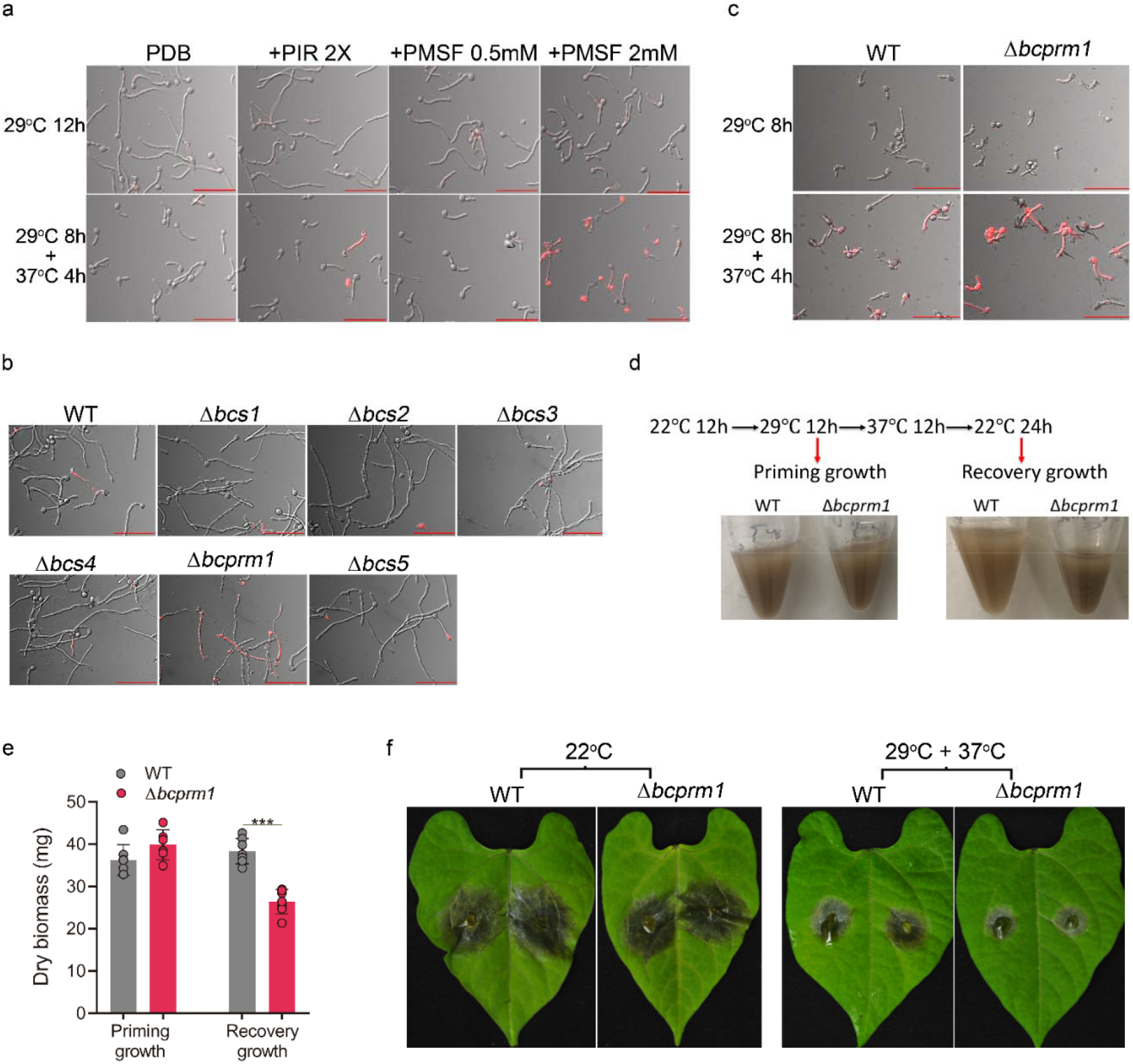
Images showing the effects of STPs on priming. **a**–**c**. Combined DIC and fluorescence (rhodamine filter) microscopic images of GTs after staining with PI (**a**,**c**) or DiBAC4(5) (**b**). Scale bars, 100 μm. **d**. Workflow of biomass measurement. **e**. Measurement of dry biomass. Graph represents at least six biological replications with overlaid individual data points. Values are presented as the mean of replicates ± s.d. Statistical differences were determined according to unpaired two-tailed Student’ t-test (\*\*\**P* < 0.001). **f**. Representative photographs of *P. vulgaris* leaves showing the effects of *bcprm1* deletion on priming pathogenicity under priming conditions.

